# Probing protein-protein interactions with drag flow: A case study of F-actin and tropomyosin

**DOI:** 10.1101/2025.05.14.653996

**Authors:** Camille Bagès, Morgan Chabanon, Wouter Kools, Thomas Dos Santos, Rebecca Pagès, Maria Elena Sirkia, Cécile Leduc, Anne Houdusse, Antoine Jégou, Guillaume Romet-Lemonne, Hugo Wioland

**Affiliations:** Université Paris-Cité, CNRS, Institut Jacques Monod, F-75013 Paris, France; Université Paris-Saclay, CNRS, CentraleSupélec, Laboratoire EM2C, 91190, Gif-sur-Yvette, France; Structural Motility, Institut Curie, Université Paris Sciences et Lettres, Sorbonne Université, CNRS UMR144, Paris 75248, France

## Abstract

Tropomyosin are central regulators of the actin cytoskeleton, controlling the binding and activity of the other actin binding proteins. The interaction between tropomyosin and actin is quite unique: single tropomyosin dimers bind weakly to actin filaments but get stabilised by end-to-end attachment with neighbouring tropomyosin dimers, forming clusters which wrap around the filament. Force spectroscopy is a powerful approach for studying protein-protein interactions, but classical methods which usually pull with pN forces on a single protein pair, are not well adapted to tropomyosins. Here, we propose a method in which a hydrodynamic drag force is applied directly to the proteins of interest, by imposing a controlled fluid flow inside a microfluidic chamber. The breaking of the protein bonds is directly visualised with fluorescence microscopy. Using this approach, we reveal that very low forces from 0.01 to 0.1 pN per tropomyosin dimer trigger the detachment of entire tropomyosin clusters from actin filaments. We show that the tropomyosin cluster detachment rate depends on the cytoplasmic tropomyosin isoform (Tpm1.6, 1.7, 1.8) and increases exponentially with the applied force. These observations lead us to propose a cluster detachment model which suggests that tropomyosins dynamically explore different positions over the actin filament. Our experimental setup can be used with many other cytoskeletal proteins, and we show, as a proof-of-concept, that the velocity of myosin-X motors is reduced by an opposing fluid flow. Overall, this method expands the range of protein-protein interactions that can be studied by force spectroscopy.

**Graphical Abstract:** 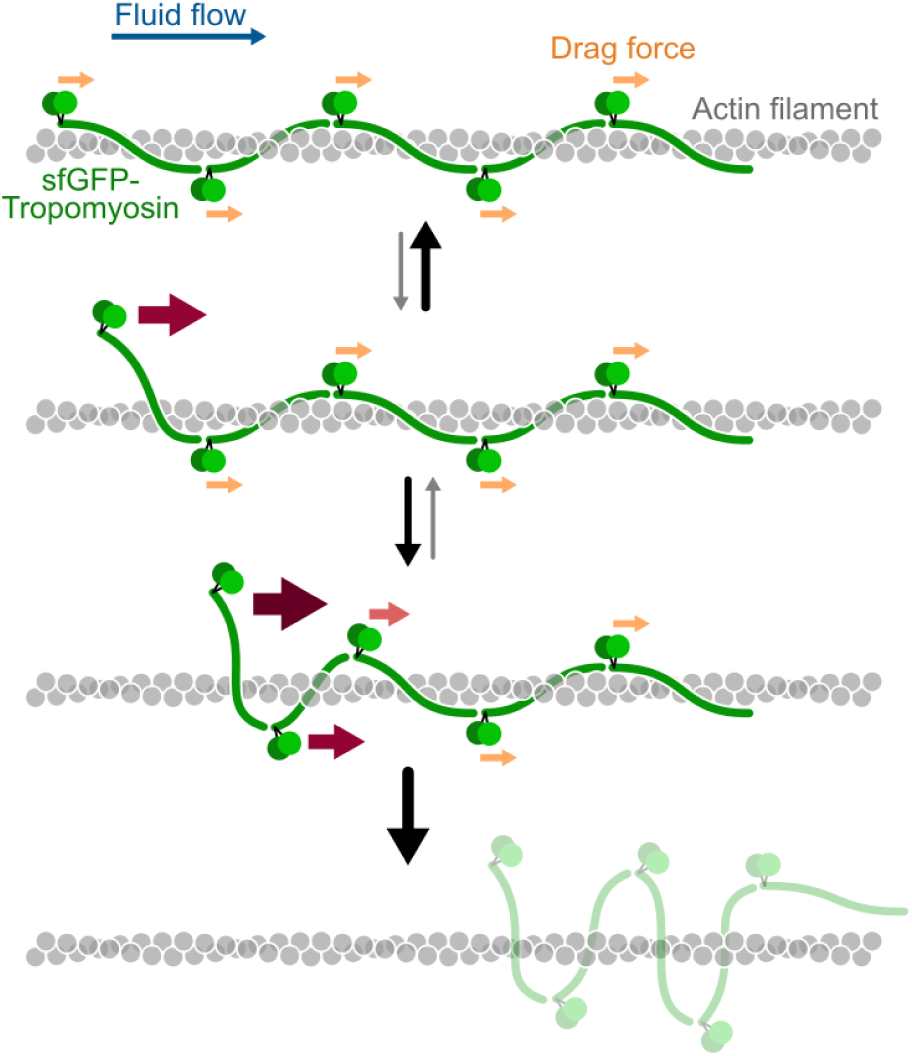

## Introduction

Tropomyosins (from thereafter, Tpm) are parallel coiled-coil dimeric proteins which bind cooperatively along actin filaments, where they control the binding and activity of other cytoskeleton proteins (Gunning et al., 2015a; Hardeman et al., 2020). As such, Tpm have been named master regulators of the actin cytoskeleton (Gunning et al., 2015a). In mammals, Tpm are expressed by four genes (TPM1-4) giving rise to dozens of isoforms by alternative splicing (Geeves et al., 2015). Tpm have first been discovered in sarcomere where, together with troponin, they regulate myosin binding and myofibril contraction. Yet, their function is not limited to sarcomeres as Tpm covers most actin filaments in various types of cells (Meiring et al., 2018). They are especially abundant in linear fiber-like networks such as stress fibers and transverse arcs (Tojkander et al., 2011). In vitro, Tpm have been shown, in an isoform-specific manner, to prevent the binding of severing protein cofilin (Christensen et al., 2017; Jansen and Goode, 2019; Ono and Ono, 2002; Wioland et al., 2021) to prevent filament bundling by fimbrin but favours that by alpha-actinin (Christensen et al., 2019, 2017), and to regulate the activity of myosin motors (Gateva et al., 2017; Reindl et al., 2022).

Among the families of actin binding proteins, Tpm have a unique way of decorating the filaments. First, each Tpm dimer binds over 6 or 7 actin subunits, depending on the Tpm isoform length. Then, while the bond between a single Tpm dimer and the filament is very transient (half-time less than 1s) (Bareja et al., 2020), Tpm dimers connect to neighbouring Tpm dimers by their N- and C-termini, to stabilize the interaction and form clusters which follow one of the two protofilaments and thus wrap around the filament (Fig. 1a) (Gunning et al., 2015b; Von Der Ecken et al., 2015). Interestingly, Tpm does not have a single binding position. In sarcomeres, Tpm adopts sequential positions leading to muscle contraction: in its relaxed state (B-state), Tpm prevents myosin binding; upon Ca2+ influx, troponin conformation change shifts Tpm laterally allowing weak myosin binding (C-state); myosins then further push Tpm away (M-state) enhancing myosin binding and motor activity (Craig and Lehman, 2001; Tobacman, 2021). Moreover, the position of cytoplasmic Tpm on the actin filament depends on both actin and Tpm isoforms (Lehman et al., 2000; Selvaraj et al., 2023). Finally, the interactions between actin filaments and Tpm are particularly difficult to study due to their weak, loose and dynamic connections, in addition to the cooperative wrapping of Tpm clusters around the filaments. As a consequence, Tpm are likely to show behaviours that one would not observe with most other actin binding proteins. It is thus worth studying the cluster kinetics in more detail, with appropriate tools.

**Figure 1:**
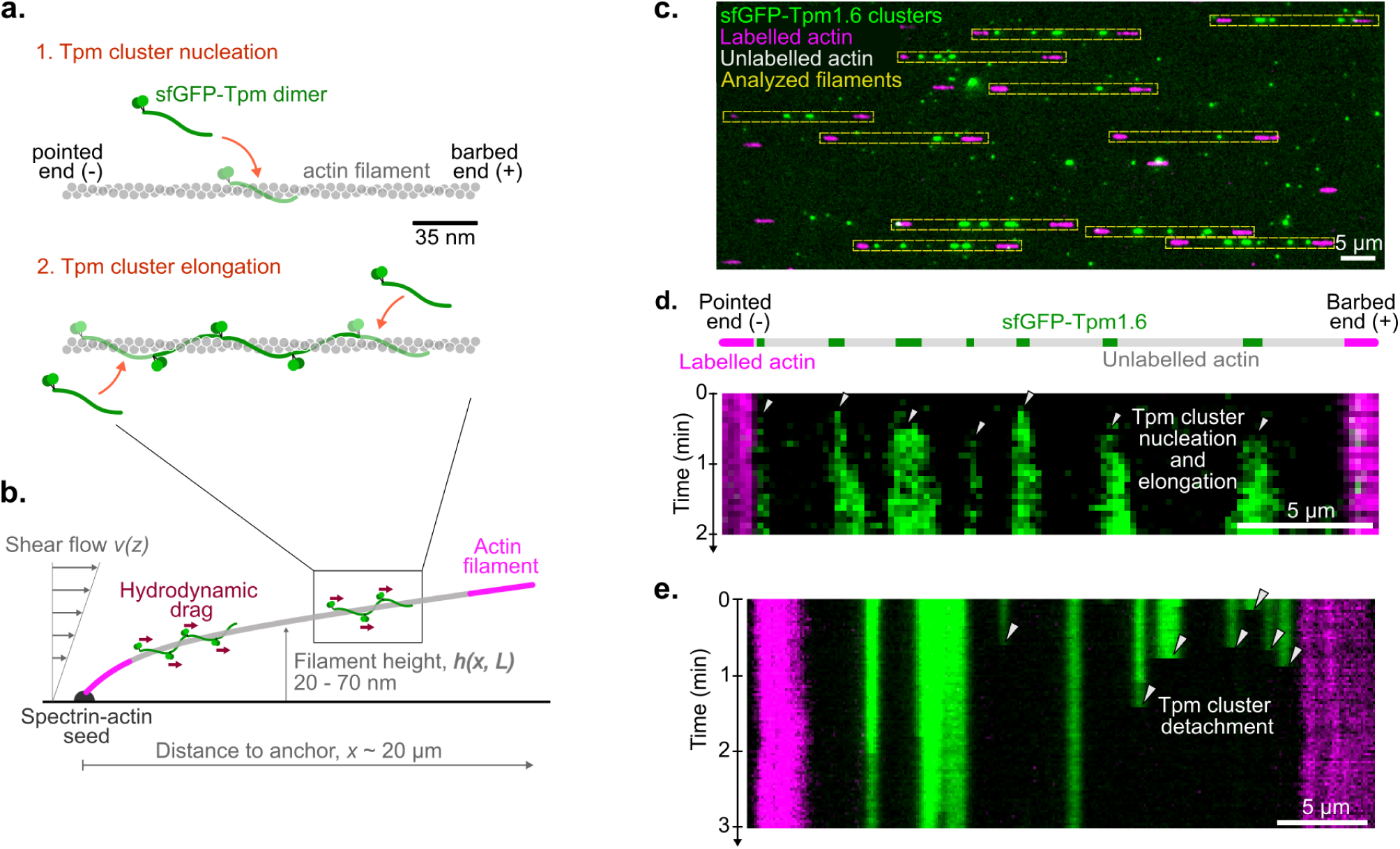
Tpm cluster assembly and detachment. a. Sketch of a Tpm cluster formation on an actin filament. The first step is the nucleation of the cluster where one Tpm dimer binds to an actin filament. Then, the cluster elongates by the addition of Tpm dimers by the interaction of their N- and C-termini, following one of the two actin protofilaments and thus wrapping around the filament. b. Experimental setup: actin filaments are polymerised from surface-anchored spectrin-actin seeds, with fluorescently labelled segments (magenta) at the two ends, and a long unlabelled middle segment (grey) over which Tpm clusters (green) are bound. All the following experiments are performed using labelled alpha skeletal actin and unlabelled beta cytoplasmic actin. Inside the chamber, the shear flow generates a hydrodynamic drag over the Tpm clusters and the actin filaments, putting the latter under tension. c. Fraction of a typical field of view showing Tpm clusters (green) bound to single actin filaments (unlabeled segment between two labeled segments shown in magenta). Scale bar 5µm d. Typical kymograph showing the nucleation of Tpm clusters (green) over a single actin filament (unlabeled segment between two labeled segments shown in magenta). e. Typical kymograph showing the detachment of Tpm clusters (green) from a single actin filament. The filament is constantly exposed to a shear flow of 13 750 /s. Note that kymographs d. and e. correspond to two different filaments.

Force spectroscopy represents a powerful approach to probe protein-protein interactions, notably to explore their force response (e.g. slip bonds vs. catch bonds) (Leckband, 2000; Neuman and Nagy, 2008; Sánchez et al., 2022) and derive cues on the energy profile (Dudko et al., 2006, 2008; Evans, 2001; Leckband, 2000). To our knowledge, force spectroscopy has not been applied to actin-tropomyosin interaction, mostly because none of the classical methods are adapted to tropomyosin. In typical experiments, each protein is anchored to a surface, which can be that of a bead, vesicle or living cell, often via a DNA linker. The proteins are put into contact to bind and pulled apart until they unbind, with a controlled force between 0.1 and 10^4^ pN, depending on the method (Sánchez et al., 2022). Classical examples of force spectroscopy instruments include optical traps, atomic force microscopy (AFM), magnetic tweezers, micropipettes, or acoustic waves (Leckband, 2000; Neuman and Nagy, 2008; Sánchez et al., 2022). Hydrodynamic drag forces have also been used to apply forces to protein-coated beads, rolling over a ligand-coated surface (Alon et al., 1995; Pierres et al., 1995), and to pull on actin filaments, thereby allowing the indirect application of forces at the formin-filament interface (Cao et al., 2018; Courtemanche et al., 2013; Jégou et al., 2013) or at the interface between the Arp2/3 complex and filaments at branch junctions (Cao et al., 2023; Ghasemi et al., 2024; Pandit et al., 2020).

However, all of the above force spectroscopy methods present limitations. They all have a finite force range and usually do not go under 0.1 pN, making them ill-suited for weak and transient protein-protein interactions. Moreover, most setups rely on protein attachments to surfaces, which might alter protein-protein interactions. Finally, these methods typically pull on a single protein pair. For proteins that assemble cooperatively onto a substrate into a complex, being able to pull simultaneously and equally on all proteins would allow probing the mechanical response of the entire complex, rather than that of a single protein within the complex. Tpm presents the additional challenge that the cluster is wrapped around the actin filament, adding a geometric constraint which maintains the Tpm cluster bound to the actin filament.

In order to circumvent these limitations, we thus designed a novel method in which a hydrodynamic drag force is applied directly to the proteins of interest by imposing a controlled flow inside a microfluidic chamber. The unbinding of the protein of interest from its surface-anchored substrate is directly visualised with fluorescence microscopy, as the protein is washed away by the flow. This approach bypasses the need for surface anchoring (apart from the anchoring of the substrate to the bottom of the microchamber, far from the interaction site). Overall this method aims at widening the range of protein-protein interactions that can be studied by force spectroscopy.

After presenting this setup (Fig. 1), we use it to apply forces that provoke the detachment of entire Tpm clusters from actin filaments (Fig. 1 and 2). We probe how salt concentration and Tpm isoforms regulate the interaction of Tpm with actin filaments (Fig. 3), we derive the detachment time as a function of the applied force (Fig. 4) and propose a cluster detachment mechanism (Fig. 5). Finally, to show that this setup can be used with very different actin binding proteins, we measure the effect of drag force on the molecular motor myosin-X (Fig. 6).

**Figure 2:**
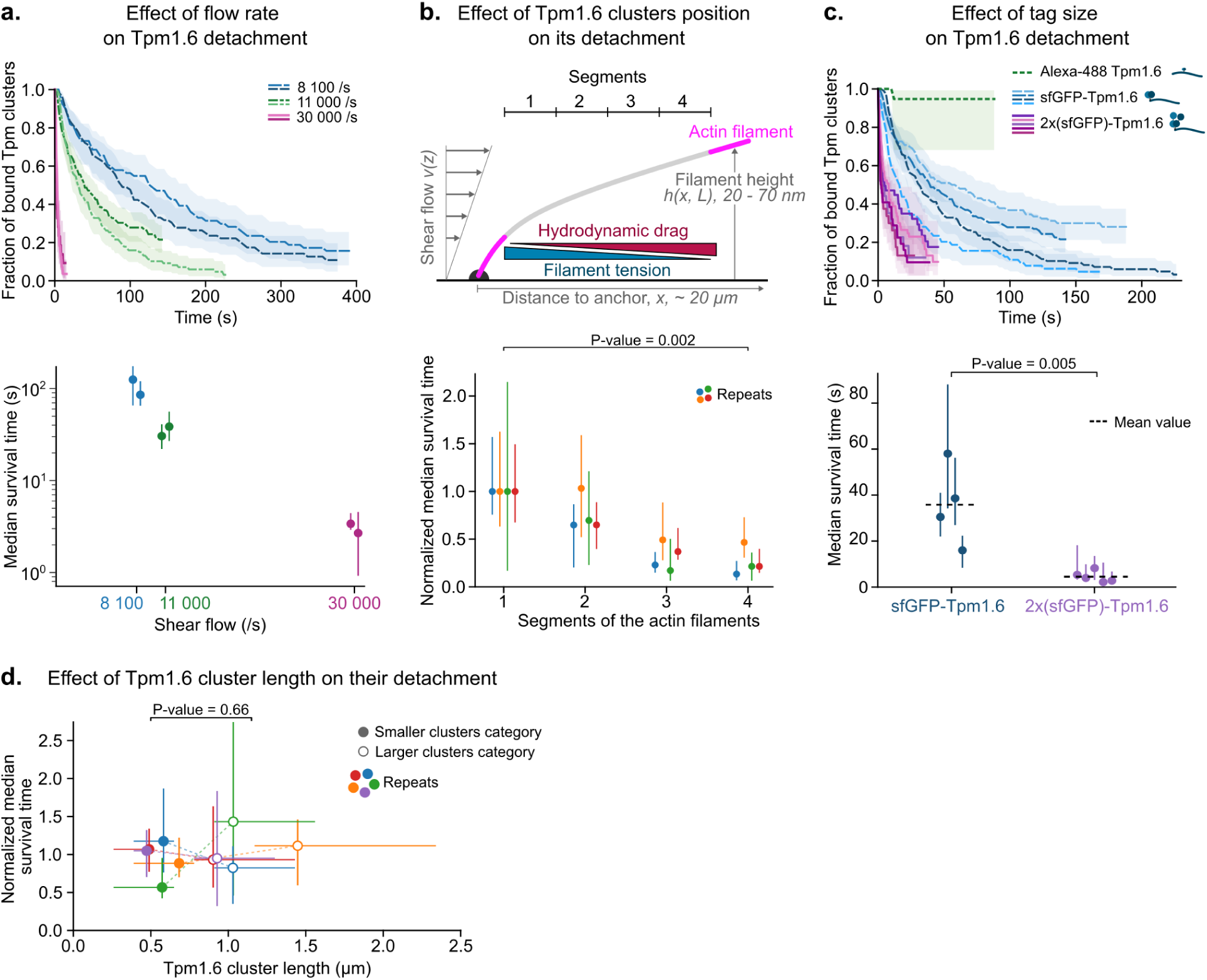
Drag force on Tpm clusters causes their detachment. a. sfGFP-Tpm1.6 cluster detachment is accelerated by the fluid flow (given as shear flow near the surface). (Top) Each curve is the survival fraction for an experiment, i.e. the evolution of the fraction of bound Tpm1.6 clusters. (Bottom) Median survival time extracted from the survival fractions, i.e. time at which 50% of the Tpm1.6 clusters have detached from the filaments. If the survival fraction does not go below 0.5, it corresponds to the median survival time extracted from the single exponential fit of the curve. Number of Tpm clusters analyzed, at 8 100 /s: N = 64, 68, 80; at 11 000 /s:, N = 140, 113; at 30 000 /s: N = 60, 25. b. sfGFP-Tpm1.6 clusters detach faster from downstream actin segments. (Top) The unlabelled part of the actin filament was split into 4 segments of equal length. The Tpm cluster population was splitted depending on the segment they belong to. (Bottom) Median survival time calculated for each Tpm cluster population, depending on the segment they are bound to, normalized by that on the first, most upstream segment. Each color corresponds to the pooled data over 1 to 3 experiments taken on the same day. Number of Tpm clusters: N = 293, 300, 107 and 318 Tpm. The shear flows were in a range 10 300 - 14 000 /s. c. Larger fluorescent tags accelerate Tpm cluster detachment. (Top) Each curve corresponds to the survival fraction of an experiment. Number of Tpm clusters for Alexa488 labelling: N = 34; single sfGFP labelling:, N = 107, 174, 140 and 78; double sfGFP labelling: N = 42, 51, 73, 46 and 58. (Bottom) Median survival times were calculated from each survival fraction curve. As only one detachment was observed during the experiment done with Alexa labelled Tpm1.6, the median survival time was not calculated. The P-value is calculated with an unpaired t-test. d. To test whether Tpm cluster length affects detachment, we compared the median survival time of the third smallest and third largest clusters. For each repeat, the median survival time is normalized by the mean of the median survival time of the two categories. One colour corresponds to the pooled data for one day of experiment with similar flow rate (± 5%). The position of each dot corresponds to the mean size of the category. Horizontal error bars represent the minimum and maximum values for each category and vertical error bars represent 95% confidence intervals. The P-value is calculated with a paired t-test over the normalized median survival time of the smaller vs larger cluster categories. There are between 1 and 3 experiments pooled with a total of N = 293, 300, 107, 318 and 148 Tpm clusters. The shear flows were in a range 8 100 - 14 000 /s. (a-c) Survival fractions: shaded areas represent 95% confidence intervals. Median survival times: error bars represent 95% confidence intervals.

**Figure 3:**
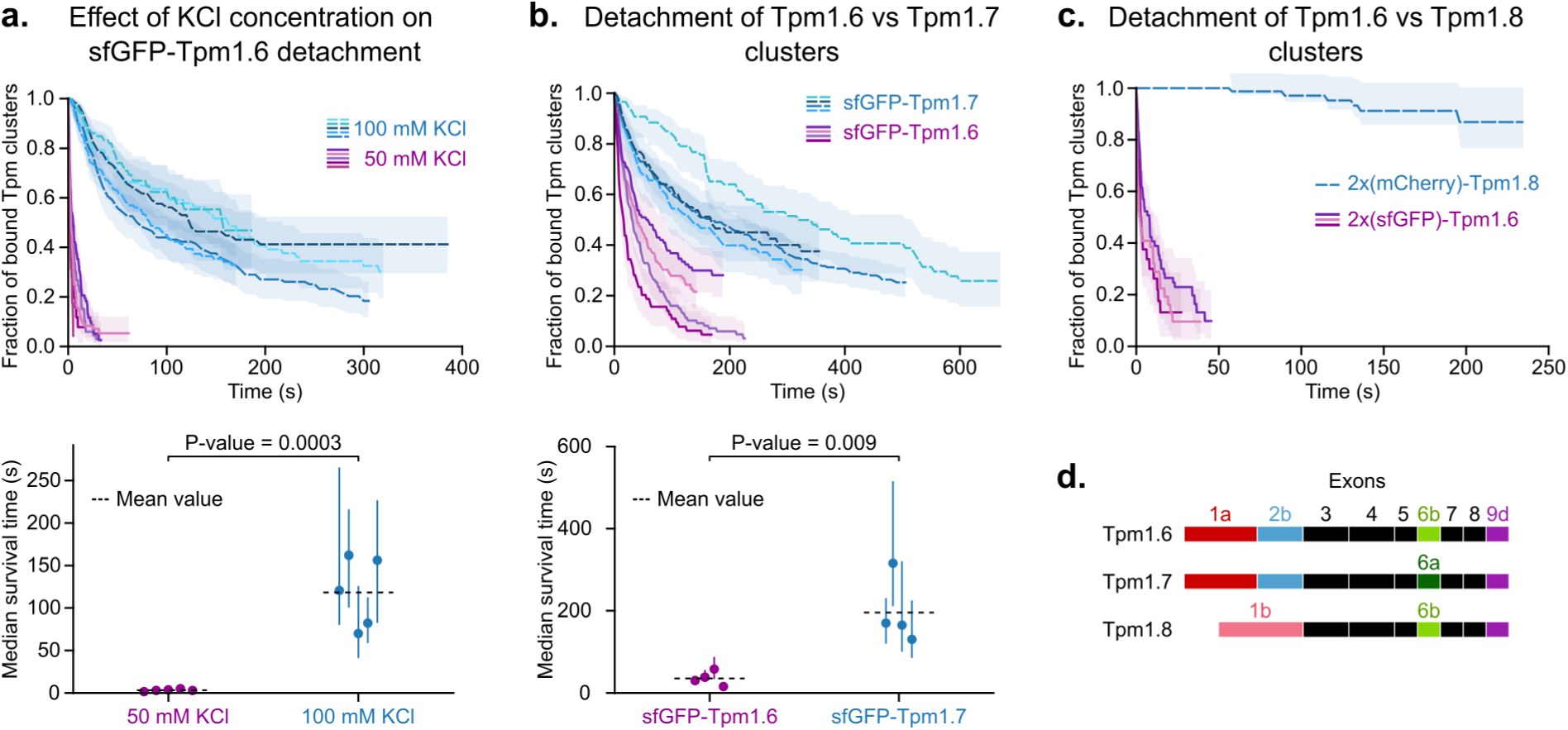
Biochemical regulation of Tpm cluster detachment. a. sfGFP-Tpm1.6 cluster detachment is accelerated by lowering the salt concentration. Survival fraction (Top) and median survival time (Bottom) of sfGFP-Tpm1.6 clusters exposed to a shear flow between 11 000 and 12 500/s. Each curve corresponds to a single experiment. Number of Tpm clusters for 50 mM KCl: N = 47, 33, 103 and 97; 100 mM KCl: N = 78, 87, 128, 89 and 170. b. sfGFP-Tpm1.6 clusters detach faster than sfGFP-Tpm1.7 clusters. Survival fraction (Top) and median survival time (Bottom) of sfGFP-Tpm1.6 and sfGFP-Tpm1.7 clusters exposed to a shear flow between 11 000 and 12 500/s. Each curve corresponds to a single experiment. Number of Tpm clusters for sfGFP-Tpm1.6: N = 140, 113, 107 and 65; sfGFP-Tpm1.7: N = 87, 238, 89 and 170. c. mCherry-mCherry-Tpm1.8 clusters detach much slower than sfGFP-sfGFP-Tpm1.6 clusters. Tpm with two tags had to be used as mCherry-Tpm1.8 clusters did not detach from actin filaments. Each curve represents the survival fraction for a single experiment. Number of Tpm clusters for 2x(sfGFP)-Tpm1.6: N = 66, 41 and 51; 2x(mCherry)-Tpm1.8: N = 92. d. Representation of the Tpm isoforms used in this study. Tpm1.6 and Tpm1.7 only differ by exon 6. Tpm1.8 is a shorter version, with exon 1b instead of 1a, and no exon 2. The fluorescent tag is fused at the N-terminus, prior to exon 1. (a-c) Survival fractions: shaded areas represent 95% confidence intervals. Median survival times: vertical error bars represent 95% confidence intervals, P-values are calculated with an unpaired t-test.

**Figure 4:**
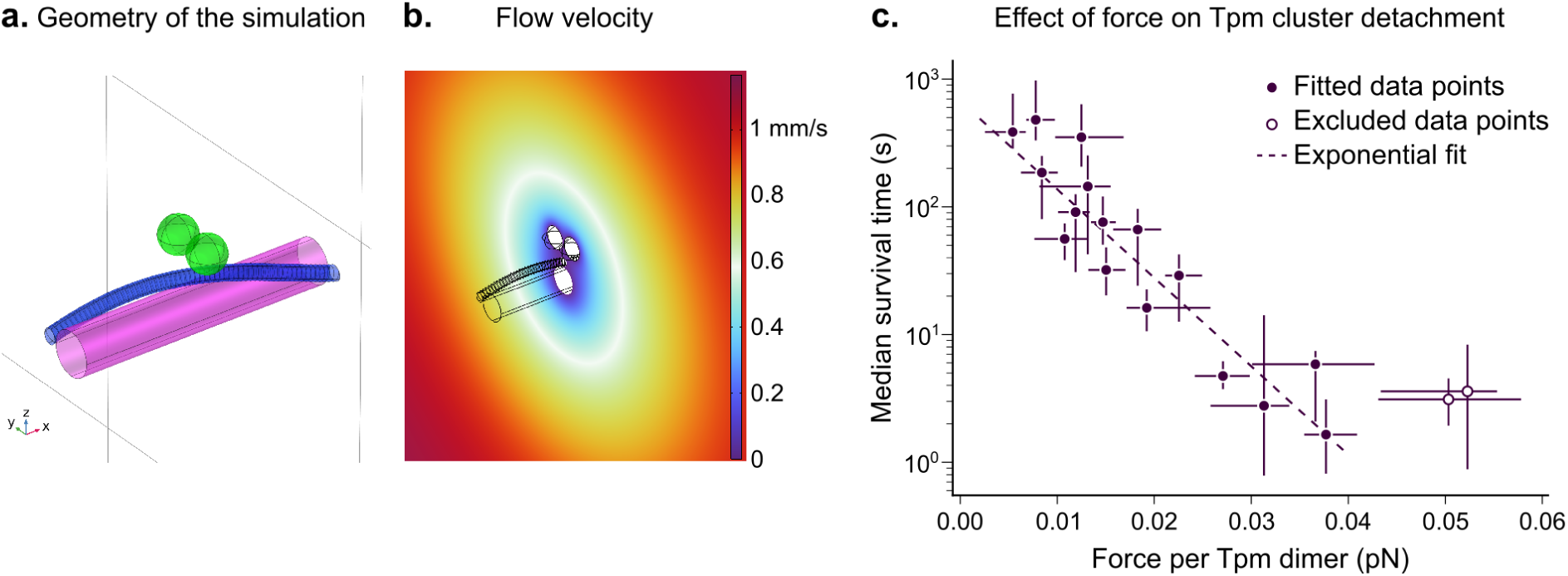
Impact of drag force on Tpm cluster detachment. a. Geometry of the sfGFP-Tpm / actin complex used in the computational fluid dynamics simulations. The magenta cylinder corresponds to a 35 nm-long segment of actin filament. The Tpm dimer is represented by a cylinder, wrapped around the filament (blue). Each sfGFP is approximated by a ball (green). Note that the simulation is run with periodic boundary conditions in the x-direction (Fig. S4a), and that the blue cylinder actually corresponds to two half-Tpm dimers, with the two sfGFPs fused at the N-terminus as in the experiments. See other configurations in Fig. S4b. b. Fluid flow velocity magnitude in the y,z plane that passes through the centre of the two sfGFPs. c. The median survival time of sfGFP-Tpm1.6 clusters decreases exponentially with the mean drag force. For each experiment the data were pooled and Tpm clusters were binned in 3 categories depending on the local flow velocity they were exposed to. For each binned population, the survival fraction was plotted and the median survival time was extracted. The position of each dot corresponds to the median force of the category. Horizontal error bars represent the minimum and maximum forces of the category and vertical error bars represent 95% confidence intervals. Total number of Tpm clusters = 826, number of experiments = 13. The dotted line corresponds to the fit with Bell’s equation (eq. 1), excluding the two largest force values (open circles).

**Figure 5:**
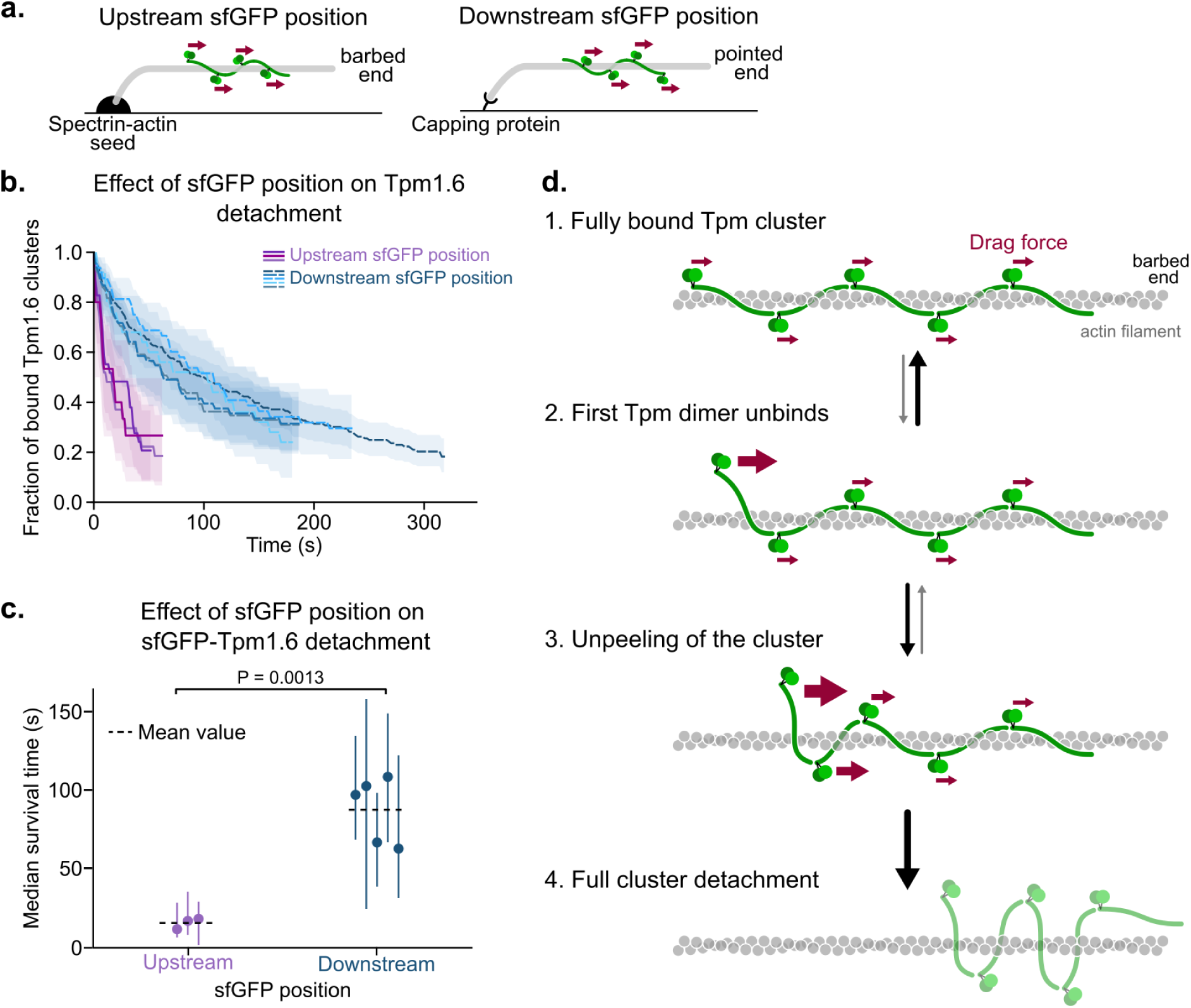
Mechanism of Tpm cluster detachment. a. To see the effect of the anchoring method of the actin filament on the Tpm clusters rate of detachment, experiments were done using spectrin actin seeds (filaments anchored at the pointed end) and other with anchored capping proteins (filaments anchored at the barbed end). b. sfGFP-Tpm1.6 clusters detach faster from pointed end anchored filaments. Detachment from unlabelled beta actin. Each curve is the survival fraction for an experiment, i.e. the evolution of the fraction of bound Tpm1.6 clusters. Number of Tpm clusters for spectrin-actin anchored filaments: N = 28, 16 and 30; capping protein anchored filaments: N = 26, 49, 162, 72 and 54.Shaded areas: 95% confidence intervals. c. Median survival time extracted from the survival fractions. Error bars represent 95% confidence intervals, P-value is calculated with an unpaired t-test. d. Model of Tpm cluster detachment. The first Tpm dimer weakly interacts with actin and is thus in equilibrium between at least two positions, bound and unbound (1 & 2). Drag force increases once the Tpm dimer is unbound (2), triggering the unbinding of the next Tpm dimer (3), further increasing the drag force until irreversibly detaching the full Tpm cluster (4).

**Figure 6:**
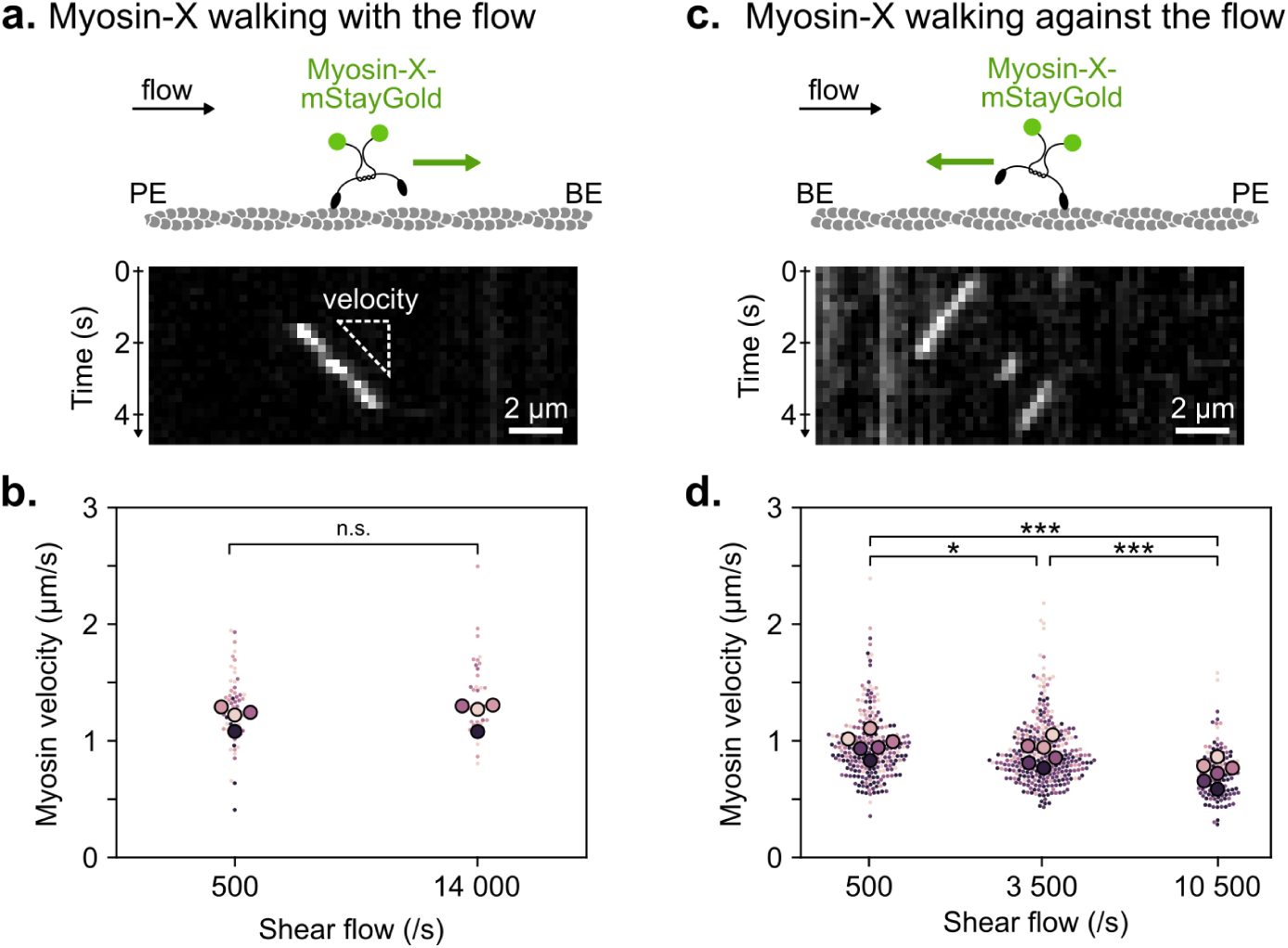
Effect of drag force on myosin-X velocity. (a, c) Sketch and kymograph of a myosin-X-mStayGold dimer walking in the same direction (a) and in the opposite direction (c) to the fluid flow. (b, d) Velocity measurements of individual myosin tracks going with (b) or against (d) the flow, exposed to different shear flow strengths. Strong shear flow significantly reduces by up to 25% the velocity of myosins going against the flow. Experiments were performed in standard F-buffer, except for the KCl concentration at 50 µM. ATP is at 0.2 mM and thanks to the constant flow is not depleted over time. Small dots correspond to individual myosin tracks, large ones to the median over a single experiment. Colours indicate independent experiments. P-values were calculated with paired t-test. P-values: n.s. = 0.11, * = 0.045, *** < 0.001. b. From slow to fast flow, N = 68 and 39 tracks, mean of median velocities: 1.21 and 1.24 µm/s, estimated average force: 0.01 and 0.13 pN. d. From slow to fast flow, N = 256, 317 and 117 tracks, mean of median velocities: 0.97, 0.90 and 0.73 µm/s, estimated average force: 0.01, 0.05 and 0.11 pN.

## Results

### Setup description

In order to visualize the dynamics of Tpm clusters onto single actin filaments under a controlled fluid flow, we use a well established setup that combines fluorescence microscopy with microfluidics (Jégou et al., 2011; Wioland et al., 2022). All experiments are performed inside a PDMS microfluidic chamber, typically 20 µm high, 800 µm wide and 1 cm long. The bottom glass surface is functionalised with spectrin-actin seeds and passivated with PLL-PEG (see Methods). The chamber is connected to three reservoirs containing different protein solutions, which are injected with a pressure controller (Fluigent MCFS). The fluid flow is measured for each inlet with flowmeters (Fluigent Flow Unit S and M).

Tpm has been shown to have a low affinity for fluorescently labelled actin filaments (Bareja et al., 2020; Schmidt et al., 2015; Wioland et al., 2021). To solve this limitation, we assemble filaments with a long unlabelled segment. This was achieved by sequentially flowing in labelled actin monomers (5 min, 0.5 µM), unlabelled actin (15 min, 1.2 µM) and again labelled actin (5 min, 0.5 µM). The assembled filaments thus contain two short fluorescent segments to reveal the two ends and a long central unlabelled segment over which Tpm binding is unaffected (Fig. 1b,c).

Tens of these assembled single actin filaments are then exposed to sfGFP-Tpm dimers (120 nM, 2 min) to form clusters on each filament (Fig. 1c,d). We mostly use Tpm1.6, which is known to bind and stabilise stress fibers (Tojkander et al., 2011). A controlled flow is then applied with fluorescent actin but no Tpm1.6 in solution, to prevent filament depolymerization, and a movie is acquired to monitor the detachment of Tpm clusters (typically 1 frame every 0.2 to 2 s).

### Observation of Tpm cluster detachment

At low flow rate, Tpm clusters remain bound, and slowly become shorter in length, meaning that Tpm dimers detach from both ends of clusters, as previously observed (Bareja et al., 2020; Wioland et al., 2021). At high flow rate, we instead discovered that entire Tpm clusters disappeared between consecutive frames: they detached as a whole and got washed away by the flow (Fig. 1e).

We tried to observe intermediate steps in the unbinding of clusters by increasing the frame rate up to 10 Hz, but even micrometer-long clusters fully disappeared between two frames. We wondered whether, once unbound, Tpm clusters carried by the flow might collide with downstream Tpm clusters and trigger their detachment. However, we never observed Tpm clusters bond to the same actin filament to detach simultaneously. Also, we never observe clusters to rebind downstream. On rare occasions (less than 1% of all clusters), some clusters seemed to slide along filaments, irregularly slipping between two frames over 0.5 to 1 µm. Given the scarcity and irreproducibility of the behaviour, we decided not to investigate it further.

In the following experiments, we quantified the rate of Tpm cluster detachment by computing their survival fraction (Fig. 2a), i.e. the fraction of clusters still bound over time, using the Kaplan-Meier method (Kaplan and Meier, 1958). Clusters that were lost, for example because the filament broke during the acquisition, were considered as censoring events (Methods). For each condition, we extracted the median survival time, i.e. the time at which half the clusters had detached.

The disassembly of Tpm clusters has always been described to happen at their ends, by the removal of single Tpm dimers (Bareja et al., 2020; Schmidt et al., 2015; Wioland et al., 2021). Since Tpm clusters are wrapped around the actin filament, the detachment of whole Tpm clusters has not been predicted or reported before, to our knowledge. Thus, to better understand Tpm cluster dynamics, we investigated what could be the cause and mechanism.

### Hydrodynamic drag is the driving force for Tpm cluster detachment

To better characterize Tpm cluster detachment, we first applied different flow rates inside the microfluidic channel. The fluid velocity follows a parabolic profile from the bottom glass coverslip to the PDMS ceiling. Close to the surface, the flow profile can be approximated by a linear velocity gradient, described as a shear flow (in units s^-1^) in the rest of the manuscript (Jégou et al., 2013). We thus compared the detachment of Tpm clusters at shear flows from 8100 s^-1^ to 30000 s^-1^ and found that the faster the flow, the faster the unbinding (Fig. 2a). Note that in all experiments, Tpm clusters are uniformly distributed along the actin filament length (Fig. S1) and that filament length for each condition is, on average, the same (around 20 µm, standard deviation 2 µm).

While the fluid flow creates a hydrodynamic drag over Tpm clusters, the friction also applies onto the actin and thus puts filaments under tension (Courtemanche et al., 2013; Jégou et al., 2013; Wioland et al., 2019). In order to tell how much the drag on the clusters and the tension of the filament each contribute to cluster detachment, we analysed the detachment depending on Tpm position along the filament.

Since filaments are attached by one end to the surface, tension accumulates from the filament free end up to the anchored end. In other words, the upstream segments are under higher tension than the downstream ones (Jégou et al., 2013; Wioland et al., 2019) (Fig. 2b top). Additionally, the height of the filament increases from the anchored end to the free end (Wioland et al., 2019) (and Fig. S2). Since the fluid velocity increases linearly from the surface, filament tension and hydrodynamic drag present opposite gradients, of which we took advantage to disentangle the relative contributions of drag force on Tpm and filament tension. (Fig. 2b top).

Within a single experiment, we then split the Tpm cluster population into four categories depending on their position along the filaments (Fig. 2b). Upstream clusters are bound to actin segments under a higher tension, while the downstream ones are exposed to a stronger drag. We systematically found that the downstream clusters detached faster, indicating that the drag over the Tpm clusters was the prime contributor to cluster unbinding (Fig. 2b bottom).

To further confirm this result, we sought to modify the hydrodynamic drag of the Tpm, without changing the shear flow. In most experiments, we use a Tpm construct fused with a sfGFP (i.e. two sfGFP per Tpm dimer) which contributes significantly to the hydrodynamic drag. In order to increase or decrease the drag force on the Tpm, we also performed experiments with 2x(sfGFP)-Tpm and Alexa488-labelled Tpm, respectively. We found that, for a given flow rate, the detachment was slowest for Alexa-Tpm (weakest drag force) and fastest with 2x(sfGFP)-Tpm (strongest drag force, median survival time about 8 fold shorter than with single sfGFP, Fig. 2c).

We also measured the detachment rate depending on the Tpm cluster length. We compared the smallest third and the largest third of the cluster population and found that longer clusters unbind as fast as shorter ones (Fig. 2d). This can be simply understood by considering that both the drag force and the Tpm-actin interaction energy scale linearly with the cluster length.

Overall, these experiments show that Tpm clusters detach by the action of the hydrodynamic drag on these clusters and that detachment does not depend on filament tension.

### Regulation of Tpm-actin interaction by biochemical factors

A number of biochemical factors can regulate Tpm-actin affinity. Here, we explore how KCl concentration and Tpm isoform affect the cluster detachment rate. The assembly of Tpm dimers, their binding to actin filaments, and the N- and C-termini locking between dimers within a cluster all rely on numerous hydrophobic and electrostatic interactions (Lorenz et al., 1995; Von Der Ecken et al., 2015). As a consequence, they are highly sensitive to salt concentration. We compared the affinity of Tpm1.6 for actin filaments at 50 and 100 mM KCl and found that clusters detach 36 times faster at the low salt concentration (Fig. 3a).

This observation is consistent with previous measurements which estimated the affinity based on the amount of Tpm bound to actin filaments in bulk solution (Eaton et al., 1975) and on the Tpm cluster assembly rate on single filaments (Schmidt et al., 2015), and found that the affinity was strongest around 100 mM KCl, compared with lower salt concentrations. While mammals contain four different Tpm genes (named TPM1 to 4), dozens of isoforms are formed by alternative splicing (Geeves et al., 2015). For a given gene, 4 out of 9 exons can vary between isoforms. Some isoforms only vary by a single exon, while others can have different N- and C-termini and have different lengths (long Tpm are composed of 9 exons and bind 7 actin subunits, while short ones lack exon 2 and bind 6 actin subunits). As a consequence, the affinity of Tpm isoforms to actin filaments can vary drastically, with an effective K_D_ ranging from 0.1 to 2 µM (Carman et al., 2021; Gateva et al., 2017; Janco et al., 2016). We first compared the median survival time of Tpm1.6 and Tpm1.7 clusters, which only differ by central exon 6 (Fig. 3d). We found that Tpm1.7 detaches 5.5 times slower than Tpm1.6 (Fig. 3b), consistent with the observation that Tpm1.7 nucleates faster than Tpm1.6 (Fig. S3). We then compared Tpm1.6 to Tpm1.8 which differ by their N-terminus (exon 1a and 2b for Tpm1.6 and exon 1b for Tpm1.8). We found that the nucleation of Tpm1.8 clusters is much faster than that of Tpm1.6 (Fig. S3). In our setup, we did not observe any detachment of mCherry-Tpm1.8 clusters. We increased the drag force by adding a second mCherry and found that mCherry-mCherry-Tpm1.8 barely detaches from filaments over several minutes (Fig. 3c), consistent with the high nucleation rate of Tpm1.8 which indicates a strong affinity for the filament.

Overall, the detachment of Tpm clusters by the hydrodynamic drag provides a new method to probe the biochemical regulation of Tpm-actin interactions. More generally, these observations show that our drag force method can detect subtle differences in protein-protein interactions. In particular, this is illustrated by the slower detachment of Tpm1.7 clusters, compared with Tpm1.6 clusters, despite a single exon difference between the two.

### Force characterization of Tpm-actin interaction

Force spectroscopy has been used to gain insight into the energy landscape of protein interactions and, overall, get information on the typical length-scales and energies of the interactions (see work by Kramer, Evans and Dudko for a complete theoretical perspective (Dudko et al., 2006, 2008)). In the simplest Bell’s model, the rate of detachment increases exponentially with the applied force:

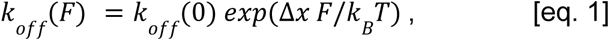

where Δ*x* denotes the characteristic interaction distance (distance between the energy minimum and barrier/maximum).

Here, one of the main challenges is to precisely estimate the hydrodynamic drag force *F* applied to Tpm clusters. Given that the detachment rate does not depend on the cluster length, and that both the drag force and the binding energy are proportional to the cluster length, we only measure the force exerted per Tpm dimer. As a first approximation, we assume the actin filament to be perfectly aligned with the flow. Being at low Reynold number, we assume the drag force *F* to be proportional to the local fluid velocity *v*, which depends on the distance *z* of the cluster to the glass coverslip. Finally the force can be calculated as *F* = *v*(*z*) · µ, were µ is the friction coefficient over the Tpm and sfGFP.

We previously measured the average conformation of actin filaments under a shear flow, i.e. the filament height *z* depending on the distance *x* from the anchored end for different flow rates and filaments lengths (Jégou et al., 2013; Wioland et al., 2019). Here, we have repeated these measurements for a larger range of shear flows and filament lengths. Of interest, at the high flow rates at which Tpm clusters detach, the height of the filaments increases linearly from ∼20 nm at the anchored end to ∼70 nm at the free end (Fig. S2).

To estimate the friction coefficient µ, we turned to computational fluid dynamics simulations (Methods). We approximated the actin filament as a cylinder, Tpm dimer as a helicoidal cylinder wrapped around the filament and each sfGFP as a single sphere (Fig. 4a, Methods). In the case of a sfGFP-Tpm dimer, we found a friction coefficient µ = 4. 0 * 10^−5^ *pN*/µ*m*. *s*. For a typical velocity *v* = 500 µ*m*/*s* (shear flow 10 000 /s), this yields a total drag force of *F* = 0. 02 *pN* on the sfGFP-Tpm dimer (0.6 pN for a 1 µm long sfGFP-Tpm cluster). Interestingly, the friction exerted on each sfGFP and on the Tpm dimer are almost equal ( µ = 1. 3 * 10^−5^ *pN*/µ*m*. *s* for one sfGFP or a Tpm dimer). As a comparison, we conducted simulations on the Tpm dimer alone (without the fused sfGFP), resulting in a friction coefficient reduced to 2. 2 * 10^−5^ *pN*/µ*m*. *s*. We also examined the effect of an addition of a second pair of sfGFP (i.e. 2x(sfGFP)-Tpm), and found an increase of the friction coefficient to 4. 8 * 10^−5^ *pN*/µ*m*. *s* (Fig. S4). For completeness, cylindrical sfGFP geometries oriented along the flow were tested, yielding slight increase in friction coefficient up to 4. 6 * 10^−5^ *pN*/µ*m*. *s* compared to spherical geometries (Fig. S4 and Table S1).

Our next goal was to calculate the cluster median survival time as a function of the applied force (per Tpm dimer). Since Tpm clusters are distributed randomly along filaments (Fig. S1), they are positioned at different heights and thus exposed to different fluid velocities. Within a single acquisition, we calculated for each Tpm cluster the time at which they departed and the force at which they were subjected (which depends on the global flow rate, the length of the filament and their position on the filament). For a given day, we then split the total cluster population into three equally-sized groups of increasing drag force. We then plotted, for each group, the survival fraction and extracted the time at which half the Tpm clusters had detached.

Results shown in Fig. 4c indicate that the cluster median survival time decreases exponentially with the applied force. It thus seems that, in first approximation, the detachment process followed the classical Bell’s model.

Surprisingly, we found that Tpm cluster detachment is extremely sensitive to force, with a median detachment time decreasing by more than 100-fold from 0.01 to 0.04 pN per Tpm dimer. This yields a typical interaction distance Δ*x* around 600 nm, an order of magnitude larger than the length for a Tpm1.6 dimer (35 nm).

We wondered whether we did not underestimate the drag force, thus overestimating Δ*x*, but uncertainty in estimating flow rates, filament height or the drag force would, at best, lead to an overestimation of the Δ*x* by a factor of 4, still yielding Δ*x* > ∼100 nm.

While Bell’s formula is well suited for single bonds whose energy potential is weakly modified by the applied force (i.e. Δ*x* * *F* smaller than the energy barrier), Tpm-actin interactions largely deviate from this simple model. Given that the interaction between a single Tpm dimer and an actin filament is very transient (half-time less than 1s) (Bareja et al., 2020), we would expect the energy barrier to be at most of the order of *k*_*B*_*T*. Instead, if we take Δ*x* = 600 *nm* and the force per Tpm dimer *F* = 0. 025 *pN*, we obtain an energy Δ*x* * *F* ∼ 4 *k*_*B*_*T*, too large to fit Bell’s model. In addition, as Tpm clusters start detaching, the drag force might increase, thus further deforming the energy landscape.

These observations lead us to propose that the detachment of Tpm clusters is a complex multi-step process.

### Model of Tpm cluster detachment

All the above experiments suggest at least two possible mechanisms: 1. All Tpm-actin bonds are considered equal and detach in a cascade effect such that the Tpm cluster slips as a whole, 2. The first Tpm dimer detaches first and progressively peels off from the cluster from the filament (Fig 5d). To decipher between the two models, we considered the position of the sfGFP pulling the cluster.

In the classical experiment (Fig. 1a), Tpm are fused with a sfGFP at their N-termini, such that the sfGFP is directly facing the incoming flow (Fig. 5a left). As a consequence, the drag force over the first pair of sfGFPs pulls directly at the very end of the Tpm cluster, favouring its peeling. In order to gain better insights into the detachment mechanism, we inverted the filament orientation by anchoring filament barbed end to a Capping Protein-functionalized surface (Fig. 5a right, Methods). Because pointed-end elongation is extremely slow, we prepolymerized filaments with a uniform but low fluorescence labelling fraction (4%, Methods). Control experiments with spectrin-actin seeds were performed with the same G-actin solution.

We found that, with both orientations, Tpm clusters detached when exposed to high flow (Fig. 5b). Yet, clusters detached 5.6 times slower from Capping Protein-anchored filaments, supporting the idea that the most upstream Tpm dimer detaches first, triggering the peeling of the full cluster (Fig. 5b,c). In the discussion and in Fig. 5d, we propose a detachment model in which the Tpm cluster unbinds in sequential steps.

### Effect of drag force on another actin binding protein: Myosin-X

Our experimental setup could potentially be used with many cytoskeletal-filament binding proteins. As a proof of concept, we tested whether the velocity of molecular motors Myosin-X could be regulated by the imposed flow.

Myosins form a large family of molecular motors that bind and move along actin filaments, or exert forces on them. In cells, myosin-X is implicated in the formation of filopodia (finger-like protrusions) where they transport Mena/VASP (Tokuo and Ikebe, 2004) and integrins (Zhang et al., 2004), and accumulate at the filopodium tip (Berg and Cheney, 2002; Popović et al., 2023). At the molecular scale, myosin-X are understood to be dimers (Berg and Cheney, 2002; Tokuo and Ikebe, 2004) whose two heads step successively on the actin filament, walking processively towards the filament barbed end (Fig. 6 a,c) (Ricca and Rock, 2010; Sun et al., 2010).

Landmark experiments on myosin-V (Clemen et al., 2005; Uemura et al., 2004) and myosin-VI (Altman et al., 2004) have shown that opposing forces in the pN-range, applied by optical tweezers, could slow down and even arrest the motor movement. On the contrary, forward forces very mildly accelerate myosin motion. We wanted to confirm this effect with our hydrodynamic drag force method, using myosin-X. We thus prepared two types of chambers. The first ones are functionalized with spectrin-actin seeds, such that actin filaments are anchored by their pointed end, and myosin-X moves in the flow direction, towards the free barbed end (Fig. 6a). In the second one, prepolymerized filaments are captured at their barbed end by surface-anchored Capping Protein, such that myosin-X moves against the flow (Fig. 6c). Filaments were then exposed to fluorescently-labelled myosin-X-mStayGold at different shear flow rates, and we quantified the velocity of single fluorescent myosin spots (see Methods).

We first compared the velocity of myosin-X moving with the flow (spectin-anchored filaments) and saw no significant difference between slow and fast shear flows (Fig. 6b). On the contrary, the velocity of myosin-X moving against the flow was reduced by increasing the shear flow: compared with 500 /s, shear flow of 3 500 and 10 500 /s decreases the velocity by 8% and 25%, respectively (Fig. 6d).

To get a rough estimate of the applied force, we used the Stoke’s law for a sphere in a viscous fluid (low Reynold number) *F* = 6π. η. *r*. *v*, where η = 10 ^−9^ *pN*/µ*m*. *s* is the water viscosity, *r* the radius of the object and *v* the fluid velocity. Myosin-X dimers are slender objects, about 35 nm-long, to which we fused two mStayGold (about 5 nm in diameter each). As an approximation, we took *r* ≈ 10 *nm* to encompass the two mStayGolds and the core of the myosin. Given that actin filaments are about 50 nm away from the surface and for a shear flow of 10 000 /s, *v* = 500 µ*m*/*s*. Overall, we obtain *F*_*myosin*−*X*_ ≈ 0. 1 *pN*. However, we did not feel confident to calculate a drag force for each individual myosin-X dimer and compare populations within a single experiment, as with Tpm (Fig. 2b). Indeed, myosins tend to make actin filaments stick to the surface, making it difficult to estimate the filament height and thus the exact velocity near myosins.

Nonetheless, these new observations are in line with early measurements on myosin-V and myosin-VI (Altman et al., 2004; Clemen et al., 2005; Uemura et al., 2004), and show that our experimental setup can be used to further probe the mechanical response of molecular motors and other actin binding proteins. Here, the main advantage compared with optical tweezers, which are usually used with myosins (Altman et al., 2004; Clemen et al., 2005; Uemura et al., 2004), is that we can parallelize the measurements: tens to hundreds of myosin tracks can be obtained in a few seconds. One current limitation is the low force range we can apply. This could be circumvented, using our setup, by increasing the fluid velocity or viscosity, or anchoring filaments to micron-sized beads (see discussion).

## Discussion

Here, we have shown that weak hydrodynamic drag forces (<0.1 pN per Tpm dimer) are sufficient to detach Tpm clusters, regardless of their length, with the detachment time decreasing exponentially with the applied force. To our knowledge, Tpm clusters have always been described to disassemble by the unbinding of individual Tpm dimers at the clusters end, and we present here the first observation of entire Tpm clusters detaching at once. The fact that clusters can easily be peeled off by weak forces might shed light on Tpm dynamics: as discussed below, this suggest that Tpm dimers dynamically adopt different positions over the actin filament and that Tpm-Tpm longitudinal bonds are longer-lived than Tpm-F-actin bonds.

Based on our force spectroscopy results and previous in vitro observations, we propose the following detachment model (Fig. 5d): the first Tpm dimer, while remaining attached to the second Tpm dimer, is in equilibrium between at least two positions, on and off the filament. The first effect of the drag force is to promote the unbound state. Away from the filament, the fluid velocity is higher and so the drag force increases. Now, the second Tpm dimer is pulled off by the drag force directly applied to it and that applied to the hanging first Tpm dimer, leading to the very rapid unbinding of the second Tpm. This process repeats on the subsequent Tpm dimers until the entire cluster peels off.

In this model, neighbouring Tpm dimers must remain bound by their N- and C-termini at least until they have detached from the filament. We suppose that the peeling is faster than the Tpm-Tpm unbinding. This model also assumes that the position of the first Tpm dimer is quite dynamic. This is supported by the low affinity of single Tpm dimers for actin (Bareja et al., 2020) and the fact that Tpm isoforms can adopt different positions over actin filaments (Craig and Lehman, 2001; Lehman et al., 2000; Selvaraj et al., 2023; Tobacman, 2021).

In a cellular context, whether Tpm clusters could detach from actin filaments, due to a mechanical pulling force, is unclear. On the one hand, fluid flows are expected to be much slower than the typical 500 µm/s we impose in vitro (for instance actin retrograde flow in lamellipodia is on the order of 10-100 nm/s (Yamashiro et al., 2023)). On the other hand, the very strong viscosity of the cytoplasm could compensate for the slow fluid flows. Likewise, friction with cellular components, including other actin filaments might induce Tpm displacement or detachment. While no protein is known to directly bind and pull on Tpm, Tpm are known to be displaced on the filaments by troponin upon Ca2+ influx, and by the binding of myosin motors (Craig and Lehman, 2001; Tobacman, 2021).

In addition, using hydrodynamic drag to probe protein-protein interactions, and in particular the detachment of Tpm clusters from actin filaments, is an original approach. This setup presents its own specificities, disadvantages and advantages compared with the many other experimental setups (optical and magnetic traps, AFM, etc.).

One main difference of our approach lies in the amplitude of the forces that can be applied. Forces tend to be weaker (less than 0.1 pN per Tpm dimer) than with other methods, and thus are well suited for weak protein-protein interactions. The force could be increased by either accelerating the fluid flow, or by holding the filament further away from the bottom surface. To do so, filaments could be attached to micron-sized beads or pedestals (Chikireddy et al., 2024).

One main disadvantage of the method is that force measurement is not as accurate as that with optical tweezers or AFM. It relies on estimates of the friction coefficient and estimates of the cluster height in the microchamber. Finally, due to thermal fluctuations, the cluster height fluctuates around the mean measured value, and so does the drag force. One solution could be to attach filaments to the surface along their length, with the caveat that the fluid velocity and thus the drag force would be reduced.

On the other hand, the use of the hydrodynamic drag presents many advantages. The setup is in itself rather simple and easy to install. First, microfluidics is now well established in many labs. Then, there is no need for complex surface/bead functionalization and proteins do not need to be attached through DNA linkers. Most importantly, when considering proteins that assemble cooperatively, the same force is applied simultaneously and equally to all proteins of a cluster. Finally, we are able to visualize the detachments of tens to hundreds of Tpm clusters in a single field of view, each cluster being exposed to a different drag force.

Hydrodynamic drag has already been used in force spectroscopy experiments. In the earlier setups, the force was applied by shear flow onto a functionalized bead or cell, rolling over a complementarily functionalized surface (Alon et al., 1995; Pierres et al., 1995). One of the main disadvantages of the method is that each bead adheres through dozens of protein-ligands complexes simultaneously such that the strength of a single bond has to be inferred from estimates of the protein density. We and other groups have also used a microfluidic setup similar to that presented in this article to apply a friction force on single actin filaments and probe the mechanosensitivity of binding proteins. It was first shown that pulling on the end-binding protein formin accelerates filament polymerization but also the unbinding of formins (Cao et al., 2018; Courtemanche et al., 2013; Jégou et al., 2013). Similarly, pulling on an Arp2/3-generated branch filament accelerates its detachment from the mother filament (Cao et al., 2023; Ghasemi et al., 2024; Pandit et al., 2020). Contrary to the setup described here, the force was not applied directly onto the protein of interest (formin, Arp2/3 complex) but on the actin filament they were attached to, a strategy which can be applied to a very limited set of proteins.

Overall, the novel setup presented here expands the range of protein-protein interactions that can be studied by force spectroscopy. Hydrodynamic drag force also presents an opportunity to study protein mechanosensitivity, as illustrated by the slowing down of myosin-X against a load (Fig. 6d). A natural expansion of this work would be to more precisely compute the applied force, increase the force range, and test other myosin constructs. While our method is not as accurate as optical tweezers, its high throughput allows us to rapidly test different conditions (myosin isoforms and mutants, biochemical conditions) and describe the force-velocity relationship. Finally, our setup can be easily adapted to other biological filaments, including microtubules (Schaedel et al., 2015), intermediate filaments and DNA (Prasad et al., 2007).

## Author contribution

CB performed and analysed all Tpm experiments, and wrote the manuscript. MC designed and ran fluid dynamics simulations. WK performed and analysed myosin-X experiments. TDS and RP produced and designed all proteins except for myosin-X. MES and AH designed and purified the myosin-X construct. CL, AJ and GRL designed the project and wrote the manuscript. HW designed and supervised the project, performed preliminary experiments and wrote the manuscript. All authors proofread the manuscript.

## Material and Methods

### Proteins and buffers

Skeletal muscle actin (UniProt P68135) was purified from rabbit muscle acetone powder following the protocol from (Spudich and Watt, 1971)] as described in (Wioland et al., 2017). Recombinant cytoplasmic β-actin (UniProt P60709) was expressed in *Pichia pastoris*, with N-terminal acetylation and His73 methylation, following the protocol described in (Hatano et al., 2020). The *Pichia pastoris* strain, expressing both NAA80 acetyltransferase and SETD3 methyltransferase, was a gift from the Balasubramaniam laboratory.

Spectrin-actin seeds from human red blood cells were purified as described in (Wioland et al., 2017). Recombinant human profilin1 (UniProt P07737) was expressed and purified as described in (Wioland et al., 2017).

sfGFPTagged tropomyosins from human TPM1 gene (UniProt P09493) were expressed in *E. coli* BL21-CodonPlus(DE3) cells as described in (Gateva et al., 2017).

A truncated Myo10-mStayGold fusion construct was made using amino acids 1-938 from human Myo10 HMM (containing the motor domain, lever arm and putative dimerization region, but not the cargo binding domains). A 19-amino acid linker (SEGGSGGSGGSGGSAASAA), a GCN4 Leucine Zipper, a SnoopTag (MDAMKRGLCCVLLLCGAVFVSPSQEIHARFRRGTVTENTICKYGYLIQMSNHYECKCIEGYVLINEDTCGKKVVCDKVENSFKACDEYAYCFDLGNKNNEKQIKCMCRTEYTLTAGVCVPNVCRDKVCGKGKCIVDPANSLTHTCSCNIGTILNQNKLCDIQGDTPCSLKCAENEVCTLEGNYYTCKEDPSSNGGGNTVDQADTSYSGSGGSGKLGDIEFIKVNKEPEA), mStayGold and a FLAG-tag were added. The construct was co-expressed with calmodulin using the baculovirus Sf9 expression system. The protein was purified using FLAG affinity chromatography followed by size exclusion chromatography, and stored at −70 °C in 5mM Imidazole pH7, 75 mM KCl, 10 mM DTT, 0.1 mM MgCl_2_, 0.5 mM EGTA, 25 µM MgADP.

All experiments, except when noted, were performed in F-buffer: 10 mM Tris-HCl pH 7.4, 100 mM KCl, 1 mM MgCl2, 0.2 mM EGTA, 0.2 mM ATP, 10 mM DTT, 2 mM DABCO.

### Data acquisition

Observations were made using a Nikon Ti2 inverted microscope equipped with a 100× oil-immersion objective and a Kinetix 22 camera. The temperature of the microfluidics chamber was maintained around 25°C using a collar objective heater (Okolab). We used an azimuthal total internal reflection fluorescence (TIRF) illumination setup (iLAS2, Gataca Systems), with 488-, 561-, and 642-nm tunable lasers (maximum power 100 mW each). The manipulation and adjustment of the TIRF setup were carried out using MicroManager software (version 2.0.1).

### Microfluidic and force application

The microfluidic chambers used in this study were made with Poly Dimethyl Siloxane (PDMS) from Sylgard. They are cross-shaped with 3 inlets and 1 outlet, 23 μm high and 800 μm wide. The PDMS chambers are mounted on standard glass coverslips. To do so, glass coverslips are first cleaned with 2% glass cleaning solution (Hellmanex™ III), 2M KOH, dH2O and ethanol. When starting an experiment, a microfluidic chamber was prepared by exposing a PDMS chamber and a cleaned glass coverslip to air plasma for 30s and then binding the two together (Wioland et al., 2022).

For a typical experiment, the microfluidic chamber was passivated overnight by injecting 20 µL of 0.1 mg/mL PLL-PEG in 10 mM HEPES pH 7.4 with a pipet. The next day, the chamber was rinsed by injecting 400 µL of F-buffer with a pipet and then connected to the microfluidic system. Spectrin-actin seeds were flowed in the microfluidics chamber at 40 pM for 2 min, and got bound non-specifically to the surface.

Actin filaments were then polymerized by alternating labelled and unlabelled G-actin solutions in order to have filaments containing two short fluorescent segments to reveal the two ends, and a long central unlabelled segment. First, short labelled actin segments were polymerized by flowing in 0.5 μM 10% Alexa Fluor 568–labeled α-skeletal G-actin for 5 min. Then, long unlabelled actin segments were polymerized by flowing in a solution with 1.2 μM cytoplasmic β G-actin and 1.3 µM profilin-1 (to avoid actin spontaneous nucleation) for 15 min. The solution of 0.5 μM 10% Alexa Fluor 568–labeled α-skeletal G-actin was flowed again for 5 min. Next, the actin filaments were exposed to a solution of 120 nM sfGFP-Tpm1.6 (dimeric concentration) to grow Tpm clusters. The filaments are exposed to the Tpm solution until there are around 5 clusters per actin filament. It typically takes 2 min. For the experiments done with different Tpm isoforms or tag sizes, the concentration of the solutions were between 15 nM and 150 nM, and the injection time between 30 s and 4 min. Finally, the solution of 0.5 μM 10% Alexa Fluor 568–labeled α-skeletal G-actin was injected at high pressure, typically 150 mBar. During this step, the Tpm clusters are exposed to a high force, leading to their detachment from the actin filaments. A solution of G-actin is injected in order to avoid filaments disassembly during this Tpm detachment step. The flow rate of the solution injected in the microfluidic chamber was recorded using Fluigent Flow Units S and M. Once almost all Tpm clusters have detached from actin filaments, we make sure the actin filaments are not stuck to the surface by setting the flow to zero.

The Tpm clusters nucleation and elongation were observed over time and acquired with the acquisition rate of 1 frame every 10 s. The Tpm clusters detachment were observed over time and acquired with the acquisition rate of 1 frame every 2 s.

### Testing the influence of the sfGFP position

#### Capping protein anchored filaments

Capping proteins were biotinylated by preparing a solution with 7.6 µM of SNAPtag-capping protein, 4.4 µM of biotin-SNAP and 10mM of DTT. This solution was left on ice for 3 to 4 hours.

The microfluidic chamber was passivated overnight by injecting 20 µL of 0.1 mg/mL PLL-PEG, containing 25% of PLL-PEG-biotin, in 10 mM HEPES pH 7.4 with a pipet. The next day, the chamber was rinsed by injecting 400 µL of F-buffer with a pipet and then connected to the microfluidic system. 5 μg/mL neutravidin was then injected in the microfluidic chamber for 2 min 30 at 150 mBar and 2 additional minutes at 20 mBar before rinsing with F-buffer for 1 min at 150 mBar. Then the biotinylated capping protein solution was diluted to 0.4 µM and injected in the microfluidic chamber for 2 min at 150 mBar.

A solution of beta cytoplasmic F-actin with 4 % Alexa Fluor 568 labelled alpha skeletal actin was prepolymerized, typically, at 10 μM for 1h. As actin filaments need to be prepolymerized before being injected in the microfluidic chamber, they cannot be composed of labelled and unlabelled segments. As Tpm interaction is weaker on labelled actin, we labelled the prepolymerized filaments as little as possible. Filaments were finally injected into the chamber, binding to the capping proteins anchored to the surface. Filaments do not interact with the surface otherwise. The rest of the experiment was done as with seed anchored filaments. The filaments were exposed to a solution of 120 nM sfGFP-Tpm1.6 to grow Tpm clusters. Then, a solution of 0.5 μM 10% Alexa Fluor 568–labeled α-skeletal G-actin was injected at high pressure, typically 150 mBar, to observe Tpm detachment from the actin filaments.

#### Spectrin-actin seeds anchored filaments

As a control, experiments were done on spectrin actin seeds anchored filaments, using the same labelling method as the capping protein anchored filaments.

A microfluidic chamber was prepared, passivated and functionalized with spectrin-actin seeds as presented previously. Then, labelled actin filaments were polymerized by flowing in a 1 μM solution of beta cytoplasmic G-actin with 4 % Alexa Fluor 568 labelled alpha skeletal actin for 10 min. As before, the filaments were then exposed to a solution of 120 nM sfGFP-Tpm1.6 to grow Tpm clusters. Then, a solution of 0.5 μM 10% Alexa Fluor 568–labeled α-skeletal G-actin was injected at high pressure, to observe Tpm detachment from the actin filaments.

### Tpm detachment analysis

Image analysis was performed manually on Fiji. Filaments are selected before analysis: only filaments with their two labelled ends clearly visible and not sticking to the surface are analyzed. Then, a kymograph is constructed for each filament. From these kymographs, the width, location and time of detachment of every Tpm cluster is manually recorded.

The fraction of surviving clusters as a function of time is computed by the Kaplan-Meier method using a Python code. The error bars correspond to the 95% confidence interval. From the survival fractions the median survival times are calculated in Python. It corresponds to the time at which half the clusters had detached i.e. the survival fraction equals 0.5. If the survival fraction does not go below 0.5, the curve is fitted by a single exponential and the median survival time is calculated from the fit.

### Force computation

The force exerted on Tpm clusters corresponds to the drag force exerted by the flow on the Tpm dimers and the two sfGFP tags associated. For one Tpm dimer, this drag force is the product of the velocity of the flow at the Tpm cluster height and the friction coefficient of the dimer and its tags.

### Calculation of the local velocity force

In the microfluidic chamber, the velocity of the flow depends on the distance from the glass coverslip z, this velocity follows a parabolic profile 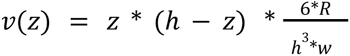 (Jégou et al., 2013). With h = 20 µm the height of the microfluidic chamber, w = 800 µm the width of the microfluidic chamber and R the global flow rate typically between 20 000 - 100 000 nL/min = 3.33 10^8^ - 1.67 10^9^ µm^3^/s.

The height of each Tpm cluster, i.e. the height of the filament at the location of the cluster, is needed to compute the local velocity of the flow. The height of filaments has been measured for filaments of 10, 20, and 30 µm (Figure S2). From these measurements, the filament height is approximated with a linear function going from 20 to 70 µm. The height of a Tpm cluster at a distance x in nm from the anchoring point of the filament is thus: 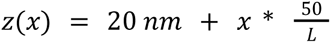 with L the total length of the actin filament in nm.

### Estimation of drag coefficient by computational fluid dynamics

Drag forces on each component of the actin-Tpm-GFP assembly were estimated by computational fluid dynamics. Geometry, meshing, computations and data visualisation were built and treated with the finite element software COMSOL Multiphysics® v.6.2 (COMSOL AB, Stockholm, Sweden).

As shown in Figure S4, the geometry was constituted of a parallelepiped fluid domain of 35 nm width and 200 nm height and depth, with the studied actin-Tpm-GFP structure at the center oriented along the width. Periodicity was assumed along the direction of the actin filament, so that the computed domain represents the length of a Tpm dimer. Actin was modeled as a cylinder of 6 nm diameter, Tpm as an helix around actin of circular base of 2 nm diameter and axial pitch 17.5 nm. sfGFP molecules were modeled as either spheres of 5 nm diameter or cylinders of 5 nm diameter and 5 nm length. sfGFP were positioned 1 nm above Tpm to account for the length of the flexible linker.

The fluid is assumed to have the properties of water. Given the nanometric dimensions of the system and the typical flow velocity of 1 mm/s, the Reynolds number is several orders of magnitude below one. Therefore an incompressible Stokes flow was solved on the fluid domain. Periodic boundary conditions on the velocity and pressure were set at the inlet and outlet, and vanishing stress boundary conditions at the other edges of the domain. No-slip boundary conditions were set at the fluid-protein surface. A mass flow rate corresponding to an average flow velocity of 1 mm/s was imposed in the direction of the actin filament.

The force applied by the fluid on each structure was evaluated by integrating the total stress tensor on the surface of the proteins. Independence of the solution with domain and mesh size was ensured for all presented data.

### Myosin-X velocity measurements

Myosin-X stock solution was thawed on the day of the experiment, centrifuged to remove aggregates (300 000 g, 20 min, 4°C) and diluted down to ∼100 nM in F-buffer with 50 mM KCl (instead of 100 mM KCl) and 0.2 mM ATP.

For experiments with myosins moving with the flow, the microfluidic chamber functionalized as for Tpm experiments. Filaments were polymerized from 1 µM rabbit α-skeletal 10% Alexa568-actin and 1 µM profilin until reaching 10-20 µm in length. With this method, filaments are anchored by their pointed end, away from which myosin-X walks.

For experiments with myosins moving against the flow, the microfluidic chamber was passivated with 0.1 mg/mL biotin-PLL-PEG (>1h, RT), rinsed with FME, functionalized with neutravidin (5 µg/mL, 5 min) and then biotin-SNAP-CP (500 nM from daily-made solution of 8 µM SNAPtag-Capping Protein, 5 µM biotin-SNAP, 10 mM DTT, left at RT for over 1h). Rabbit α-skeletal 10% Alexa568-actin filaments were prepolymerized at 10 µM for > 30 min at RT, diluted down to 0.5 µM and injected into the microfluidic chamber. Filaments would be left to bind surface-anchored Capping Protein at very low flow (<500 nL/min, shear flow <50/s) for a few minutes until the appropriate density had been reached. With this method, filaments are anchored by their barbed-end, towards which Myosin-X walks.

Myosin-X-mStayGold binding and movement was then recorded with a stream acquisition (100 to 200 ms exposure for ∼300 frames) with TIRF microscopy. For a given field of view and set of filaments, a series of movies was taken, one for each flow rate of interest. Images of filaments were taken before the myosin acquisition.

Myosin velocity was measured manually on ImageJ. For each movie, a global kymograph was generated and the user marked each individual track (segment tool). We only considered tracks that lasted at least 4 frames. Immobile myosins were excluded from the analysis. Likewise, if during a track, a myosin would stop moving, only the mobile part would be fitted.

Data was exported and plotted in Python, using panda and seaborn packages for calculating medians per repeat and plotting swarmplots. P-values were calculated with Scipy “ttest” function (with paired repeats) over the median velocities per experiment.

## Data availability

Raw data for all figures have been included as a supplementary Excel file. Further data would be available upon request to the authors.

## Supplementary information

**Figure S1:**
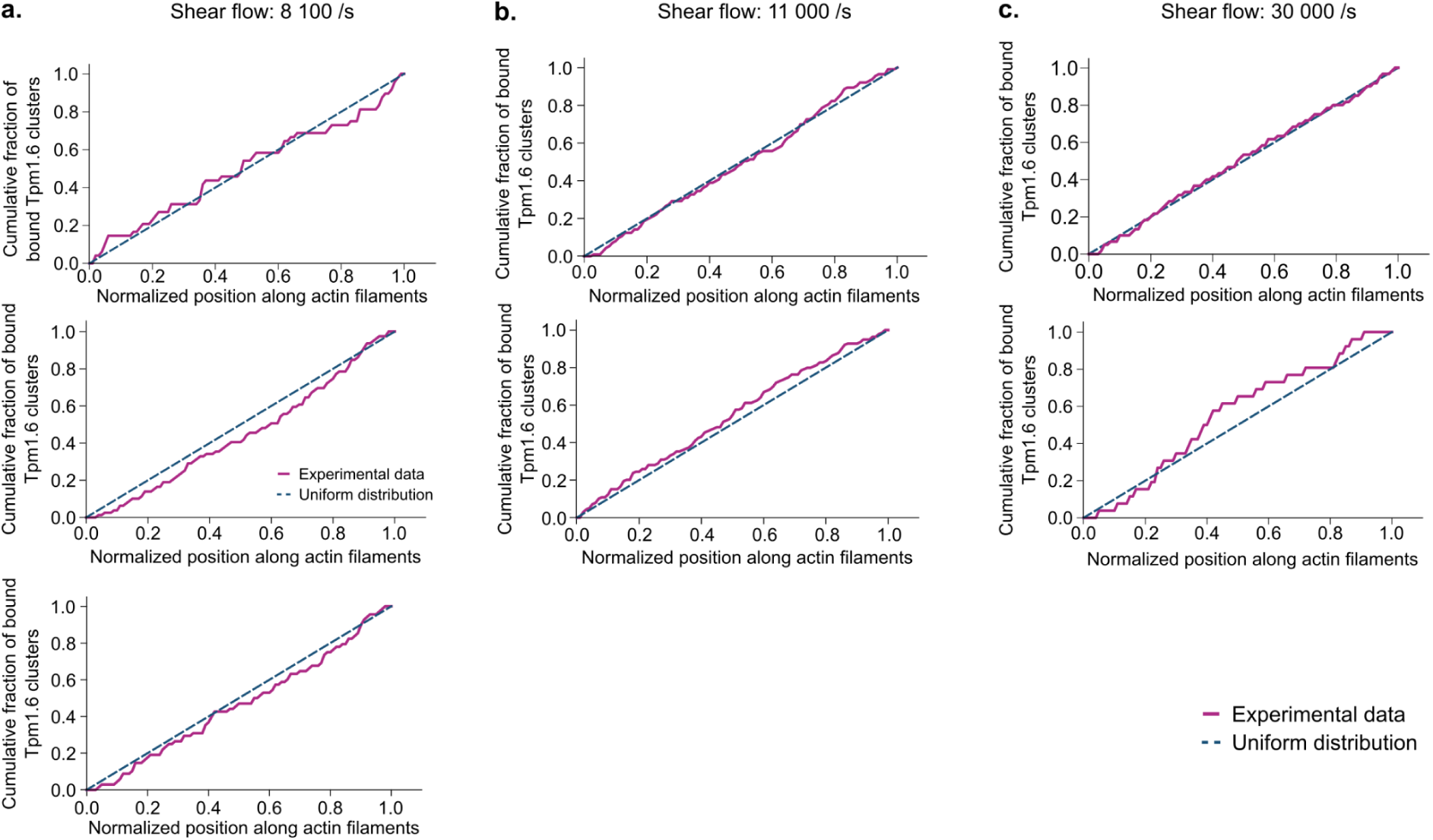
sfGFP-Tpm1.6 clusters are distributed uniformly along actin filaments, for all shear flows. Cumulative number of bound sfGFP-Tpm1.6 clusters along actin filament length. For one/a given experiment, we manually measured the position of every Tpm cluster on the unlabelled segments of all filaments. The position was normalized between 0 and 1, corresponding to the upstream and downstream ends of each unlabelled segment, respectively. We then computed the cumulative distribution of Tpm clusters, normalized by the total number of Tpm clusters. The experiments analysed here are the same as in Fig 2a. The dotted lines, “theoretical cumulative distribution”, correspond to the case where Tpm clusters are uniformly distributed along actin filaments. Number of Tpm clusters analyzed, at 8100 /s: N = 64, 68, 80; at 11000 /s:, N = 140, 113; at 30000 /s: N = 60, 25.

**Figure S2:**
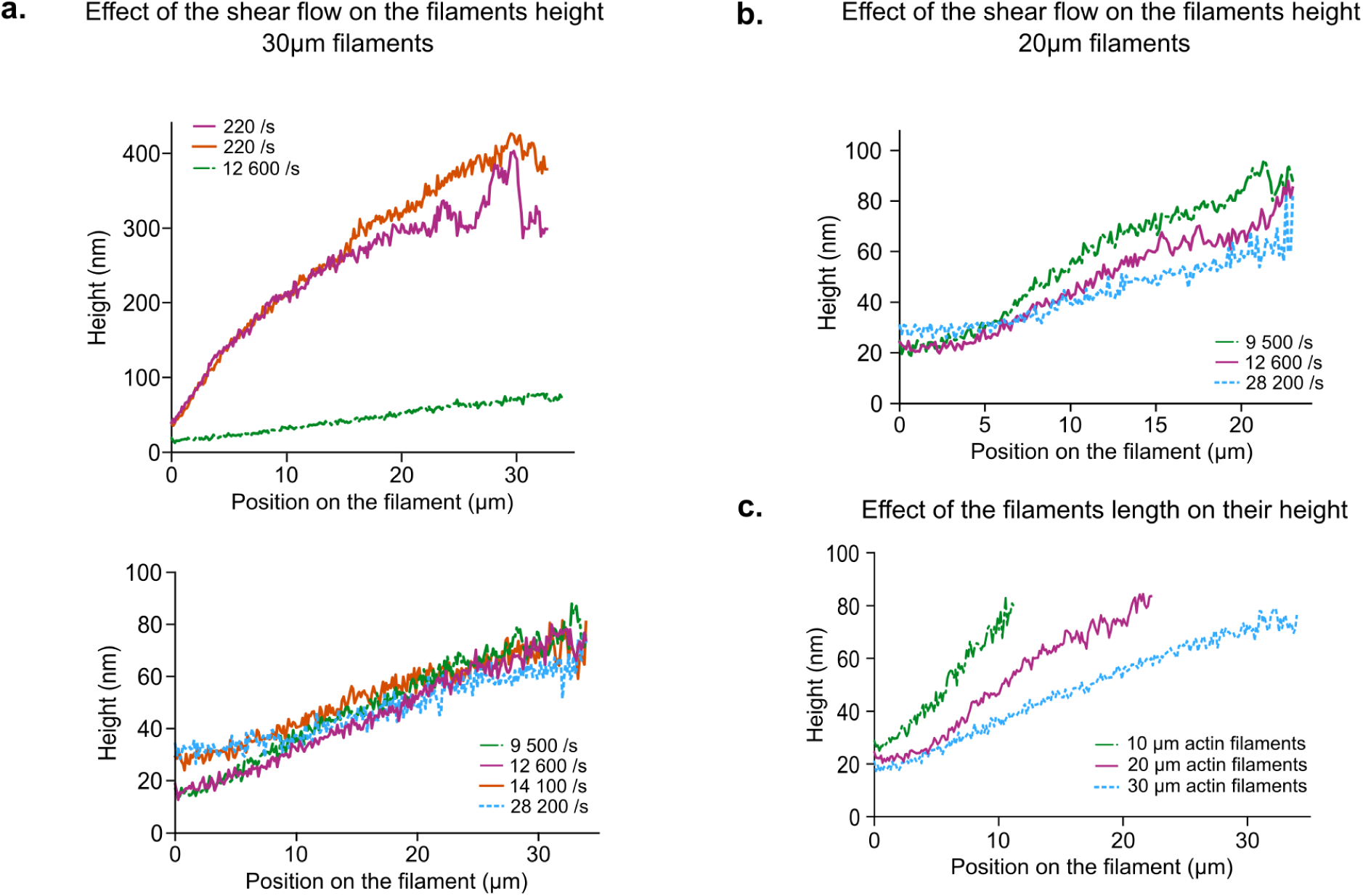
Characterisation of actin filaments height. The four graphs show the height profile of actin filaments for different shear flow and filament lengths. These height profiles were measured from actin filaments (labelled at 45.5% with Alexa-488) fluorescence intensity under TIRF illumination. Each curve was obtained by averaging the fluorescence of 10 filaments of similar length, and over 35 to 40 frames for each filament (Wioland et al., 2019). With this method, it is difficult to measure the filament height near the anchor, we thus ignored the measure in the two first pixels. In this range (0 to 0.3 μm), the filament height quickly increases from 0 to about 20 nm. a. Height profiles of actin filaments with a length around 30 µm (top) As described previously (Jégou et al., 2013; Wioland et al., 2019), at small flow gradient, here 220 /s, the filament height reaches a plateau around 300 nm from the surface. At a stronger flow rate, here 12 600 /s, the filament height appears to increase linearly. (bottom) In the range of the shear flows used in this study (8 000 - 30 000 /s), the profiles are very similar. We fitted these profiles by a linear function going from ∼ 20 nm to ∼ 70 nm. b. As observed for the ∼ 30 µm long filaments, the height profiles are very similar in the range of the shear flows used in this study. As previously, we fitted these profiles by a linear function going from ∼ 20 nm to ∼ 70 nm. c. Height profiles of actin filaments of different lengths measured at 9 500 /s. For both ∼ 10 µm, ∼ 20 µm and ∼ 30 µm long filaments, the height profiles increase linearly, from ∼ 20 nm to ∼ 70 nm. For this study every filament height profile is fitted by the same linear function: 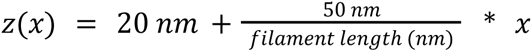 with x the position along the length of the actin filament, in nm.

**Figure S3:**
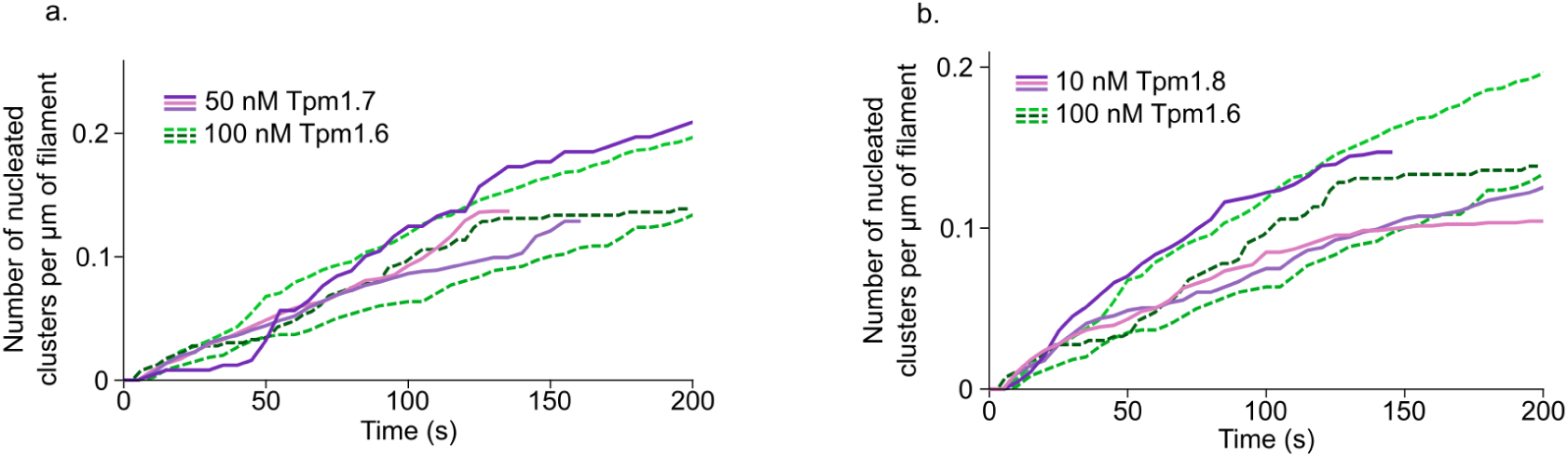
Effect of the tropomyosin isoform on the nucleation rate of clusters. Both graphs show the density of Tpm clusters over time for different Tpm isoforms and concentrations. Undecorated filaments are continuously exposed to Tpm dimers, from time 0 onwards. a. A concentration two times higher is needed for sfGFP-Tpm1.6 to nucleate clusters at the same rate as sfGFP-Tpm1.7. Number of Tpm clusters analyzed, for 50 nM sfGFP-Tpm1.7: N = 72, 183, 149; for 100 nM sfGFP-Tpm1.6: N = 55, 230, 137. b. A concentration ten times higher is needed for sfGFP-Tpm1.6 to nucleate clusters at the same rate as mCherry-Tpm1.8. Number of Tpm clusters analyzed, for 100 nM sfGFP-Tpm1.6: N = 55, 230, 137; 50 nM sfGFP-Tpm1.7: N = 72, 183, 149; 10 nM mCherry-Tpm1.8: N = 82, 108, 176.

**Figure S4:**
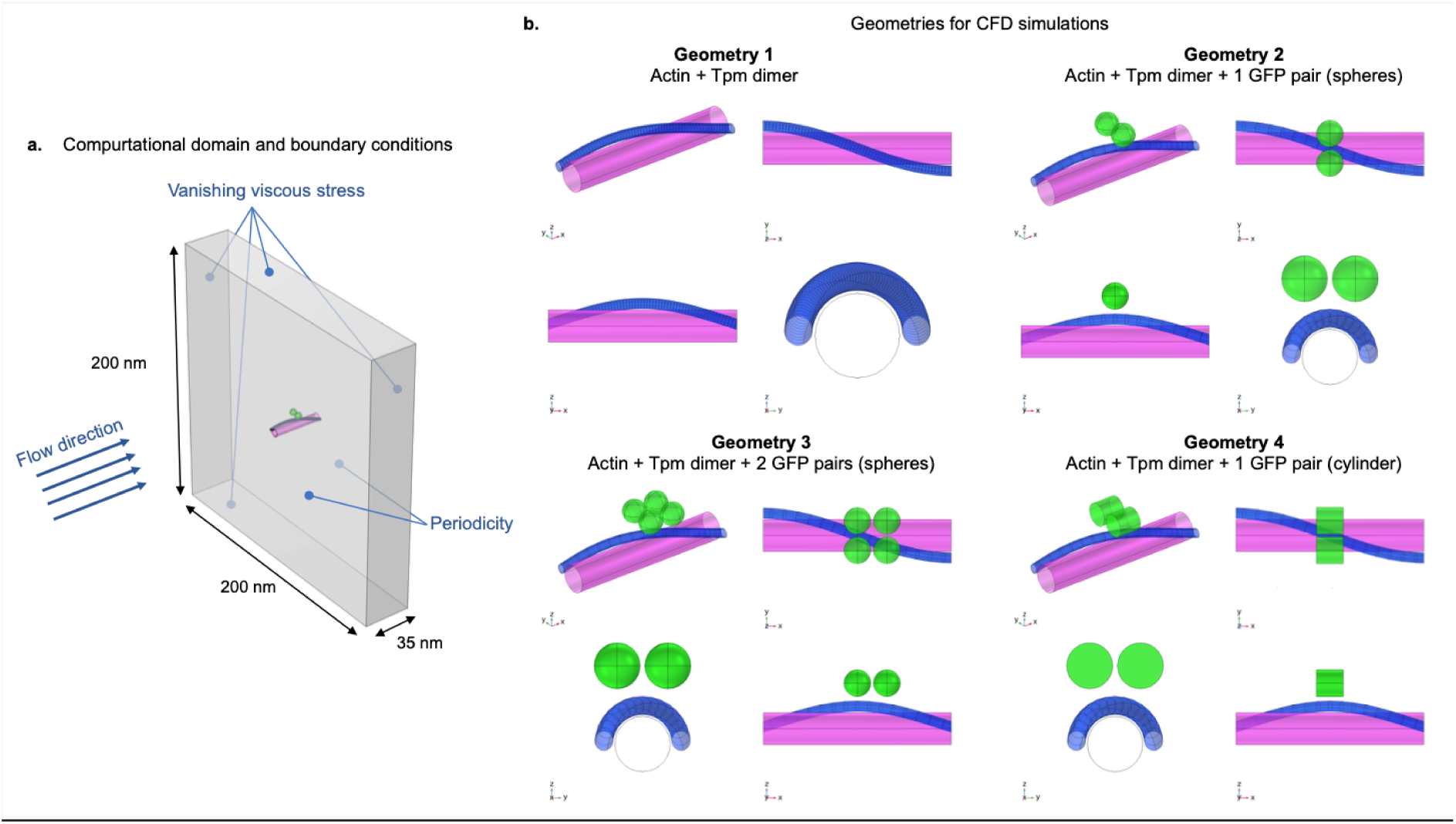
Geometries of the computational fluid dynamics simulations used to estimate the drag coefficient over sfGFP and Tpm dimers. a. Definition of computational domain and boundary conditions. b. The four geometries of Tpm dimers (blue) and sfGFP (green) constructs evaluated in this study, each with four different views.

**Table S1:** Friction coefficient obtained by computational fluid dynamics for each structure and geometry (see Fig. S4)

|  | Friction coefficient (pN/μm.s) |  |  |  |
| --- | --- | --- | --- | --- |
|  | Tpm | 1st pair of GFP | 2nd pair of GFP | Total |
| **Geometry 1**<br>**Tpm dimer** | 2.2 | x | x | **2.2** |
| **Geometry 2**<br>**sfGFP-Tpm dimer** | 1.3 | 2.7 | x | **4.0** |
| **Geometry 3**<br>**2x(sfGFP)-Tpm dimer** | 1.4 | 1.7 | 1.7 | **4.8** |
| **Geometry 4**<br>**cylindrical**<br>**sfGFP-Tpm dimer** | 1.6 | 3.0 | x | **4.6** |

